# Metformin inhibits PDGF signaling to suppress hyaluronan and IL-6 production in Thyroid Eye Disease

**DOI:** 10.1101/2025.05.19.654298

**Authors:** Farha Husain, Charkira C. Patrick, Elisa Roztocil, Steven E. Feldon, Collynn F. Woeller

**Affiliations:** Flaum Eye Institute, University of Rochester Medical Center, Rochester, New York, United States; Department of Microbiology and Immunology, University of Rochester Medical Center, Rochester, New York, United States; Center for Visual Sciences, University of Rochester Medical Center, Rochester, New York, United States; Department of Environmental Medicine, University of Rochester Medical Center, Rochester, New York, United States

**Keywords:** Thyroid eye disease, platelet-derived growth factor, hyaluronan, IL6, metformin, AMPK

## Abstract

**Background:** Thyroid eye disease (TED) is a debilitating autoimmune disorder affecting 25-50% of patients with Graves’ disease. TED is characterized by inflammation and tissue remodeling of orbital tissues. Orbital fibroblasts (OFs) and platelet-derived growth factor (PDGF) signaling promote tissue remodeling in TED. While metformin’s therapeutic potential has been proposed in various inflammatory conditions, its role in modulating PDGF signaling in TED remains unexplored.

**Methods:** OFs were isolated from TED (n= 14) and non-TED subjects (n=4). OFs were treated with PDGFβ (25 ng/mL) and/or AMPK activators metformin (1-5 mM) and AICAR (0.4-1 mM). Hyaluronan (HA) production was assessed via agarose gel electrophoresis and ELISA. Inflammatory mediators (IL6 and IL8) were measured by ELISA. Protein expression and signaling pathways were analyzed by Western blot.

**Results:** TED OFs showed enhanced HA synthesis (∼3-fold increase) and IL6 and IL8 responses to PDGFβ. PDGFβ treatment suppressed AMPK phosphorylation in a dose-dependent manner. Metformin increased AMPK phosphorylation (3.2-fold) and decreased IL6 and IL8 production. Both metformin and AICAR attenuated PDGFβ-induced HA production (56-68% reduction), IL6 (∼50% reduction), and IL8 (∼65% reduction) production in TED OFs.

**Conclusions:** This study demonstrates that PDGFβ suppresses AMPK signaling while AMPK activation by metformin counters PDGFβ-induced responses. These findings suggest that metformin is a potential therapeutic option for TED through modulation of HA and inflammatory cytokine production.

## Introduction

Thyroid eye disease (TED) represents a significant clinical challenge, causing orbital disfigurement, vision impairment, and decreased quality of life in patients with autoimmune thyroid disorders, particularly Graves’ disease (GD).^1, 2^ Affecting approximately 25-50% of patients with GD, TED has an estimated incidence of 16 cases per 100,000 women and 2.9 cases per 100,000 men annually in the general population. This prevalence substantially burdens affected individuals and healthcare systems.

The pathogenesis of TED involves several distinct but interconnected processes. TED is characterized by immune cell infiltration, tissue remodeling, expansion, and fibrosis of orbital tissues, adipogenesis, and glycosaminoglycan accumulation.^3^ Several key pathogenic features drive disease progression. For example, immune cell infiltration of ocular tissue leads to activation of orbital fibroblasts (OFs), which are key effector cells in TED. Activated OFs subsequently release pro-inflammatory cytokines such as IL6 and IL8, amplifying disease progression.^4^ Activated OFs also produce excessive extracellular matrix components, particularly collagen and the glycosaminoglycan, hyaluronan (HA). The extracellular accumulation of HA attracts inflammatory cells and water into the tissue, resulting in orbital congestion, edema, and characteristic protrusion of the eye seen in TED patients.^3,5,6^

TED is commonly treated with corticosteroids and immunosuppressive drugs to improve symptoms and shorten the disease progression.^7^ High-dose glucocorticoids show response rates of a little more than half in active TED, but their efficacy is often temporary, and they carry significant side effects. The recent introduction of teprotumumab, an anti-insulin-like growth factor-1 receptor (IGF-1R) monoclonal antibody, has demonstrated response rates of approximately 80%. However, this breakthrough therapy has significant limitations, including high cost, limited availability, and side effects such as hearing loss, muscle spasms, nausea, and hyperglycemia. Furthermore, disease recurrence after treatment remains a significant concern.^8–11^ Alternative therapies under investigation include rituximab (an anti-CD20/B-cell blocking antibody) and tocilizumab (an anti-IL6 receptor antibody), but results remain preliminary. These treatment limitations underscore the need for continued research into the underlying disease mechanisms to develop more effective, targeted, and accessible therapies.

While TED pathogenesis is not fully understood, several key molecular pathways have been implicated. Autoantibodies directed against thyroid-stimulating hormone receptor (TSHR) activate inflammatory and HA biosynthetic pathways.^12,13^ The IGF-1R pathway also promotes HA accumulation and adipogenesis.^14–16^ Platelet-derived growth factor (PDGF) receptor signaling is also elevated in disease.^17,18^ PDGF receptors show increased expression in orbital tissues from TED patients, with PDGFRβ being particularly abundant in TED OFs.^18^

PDGF cytokines comprise five different isoforms (α, β, α/ β, C, and D) that mediate their effects via binding tyrosine kinase receptors (PDGFRα and PDGFRβ). PDGFs function in part through the phosphatidylinositol-3 kinase (PI3K)/AKT signaling cascade. PI3K/AKT signaling promotes proliferation, extracellular matrix production, and secretion of inflammatory cytokines in OFs.^18–20^ Multiple studies have reported elevated levels of PDGF and PDGFRβ in orbital tissue and fibroblasts of TED patients.^18,20^ Previous work has revealed that inhibiting PDGF receptor signaling may help mitigate the progression of the disease and minimize symptoms.^20^ Thus, targeting PDGF signaling presents a potential therapeutic option for treating TED.

Despite advances in understanding the role of PDGF in TED, a significant knowledge gap exists regarding the interaction between PDGF signaling and the adenosine monophosphate-activated protein kinase (AMPK) pathway in OFs. Recent studies suggest AMPK’s involvement in regulating inflammation, adipogenesis, and HA production in TED.^21,22^ Metformin, the most widely prescribed medication for type 2 diabetes, indirectly activates AMPK by inhibiting mitochondrial Complex I.^23^ With over 60 years of clinical use, metformin has a well-established safety profile. It has shown efficacy in other autoimmune conditions, including rheumatoid arthritis and systemic lupus erythematosus.^24–27^ These properties make metformin an attractive candidate for repositioning as a TED therapy. In this study, we investigate the therapeutic potential of metformin in TED through its effects on PDGF-induced OF activation. Our approach offers several advantages over existing treatments: metformin is inexpensive, widely available, has minimal side effects, and could potentially address multiple pathogenic pathways simultaneously through AMPK activation.

## Material and methods

### Cell culture and reagents

Orbital fibroblasts (OFs) from TED patients and control subjects with no history of TED were obtained during orbital decompression surgery for severe TED or non-TED eye surgeries at the Flaum Eye Institute. Tissue procurement procedures were conducted following the Declaration of Helsinki and approved by the University of Rochester Medical School Research Subjects Review Board. Informed written consent was obtained from all subjects before surgery. Demographic data of the patients used in this study are listed in **Table 1**. OFs were cultured in Dulbecco’s Modified Eagle’s Medium/Nutrient Mixture F-12 (DMEM/F12) supplemented with 10% fetal bovine serum (FBS) and antibiotics. All media and supplements were purchased from Gibco (Carlsbad, CA) or Corning Life Sciences (Corning, NY). FBS was from Hyclone (Logan, UT). Cells were serum-starved with 0.1% FBS-DMEM/F12 medium for at least 24 hours before treatment. Cells were then treated with PDGFβ (PeproTech, Cranbury, NJ) for 72 hours to drive proliferation, HA, and cytokine production. To examine the effect of AMPK activators on the regulation of signaling pathways, cells were pretreated with metformin (MilliporeSigma, Burlington, MA, cat no. 3172405GM) or AICAR (Tocris Bioscience, Bristol, UK, cat no. 2840/50) for 1 hour before addition of PDGFβ for 72 hours.

**Table 1.** Demographic data of patients with non-TED/TED from this study.

| **Category** | **Non-TED** | **TED** |
| --- | --- | --- |
| Number | 4 | 14 |
| Females | 2 | 7 |
| Males | 2 | 7 |
| Age Range | 45-58 (54.5) | 32-74 (52.2) |
| Corticosteroids | 0 | 5 |
| Smoking Status |  |  |
| Yes | 1 | 5 |
| No | 0 | 5 |
| Former | 0 | 4 |
| Unknown | 3 | 0 |

### Determination of the HA level via agarose gel electrophoresis

To analyze the level of secreted HA, conditioned media were collected and concentrated 10-fold using a Vivaspin® 500 centrifugal concentrator (Sartorius, cat no.VS0102). Electrophoresis of HA was performed according to the method of Lee and Cowman.^28^ 50 μL of concentrated media samples were incubated with 0.5 mg/ml of Proteinase K (MilliporeSigma, cat no. P5568) at 55° C for 1 hour on a shaker. Samples were mixed with loading buffer (0.02% bromophenol blue and 2M sucrose in Tris-Borate-EDTA (TBE) buffer). HA was separated on a 1% agarose gel (SeaKem® HGT Agarose from Lonza, Basel, Switzerland) in 1X TBE buffer. Gels were stained with 0.005% Stains-All (MilliporeSigma, cat no. E9379) in 50% ethanol overnight under a light-protective cover. To destain gels, they were placed in 30% ethanol, incubated for several hours in the dark, and photographed on the Chemi-Doc MP Imaging System (Bio-Rad, Hercules, CA). 15 MDa HA (Lifecore Biomedical, Chaska, MN, HA15M-1) and Select-HA HiLadder (Echelon Biosciences, Salt Lake City, UT) were used as molecular standards. Densitometry analysis was done using Image Lab software. To confirm the staining was HA specific, aliquots of the media samples were treated with 1 U/mL of Hyaluronidase (MilliporeSigma, H1136) and incubated overnight at 37°C. The following day, samples were processed and separated on an agarose gel, as described above.

### Quantitation of HA

OFs were plated in 6-well plates, and treatment was performed for 72 hours. After each experiment, we directly collected conditioned media from the treated cells to quantify the amount of HA produced and secreted into the media. The Hyaluronan DuoSet ELISA kit was used for HA quantification (R&D Systems, Minneapolis, MN). Each experiment was performed in triplicate, and HA levels were assayed by a microplate reader (Bio-Rad, iMark™).

### Quantification of IL6 and IL8 levels

Conditioned media were collected from cell cultures following 72 hours of treatment, briefly centrifuged to remove cellular debris, and appropriately diluted. High-affinity ELISA plates were coated overnight at 4°C with capture antibodies specific for IL6 (BD Pharmingen, cat. #554543) or IL8 (Endogen, cat. #M-801). After washing with PBS containing 0.05% Tween-20 (PBST), plates were blocked with 1% BSA in PBS for 1 hour at room temperature and rewashed. Diluted conditioned media samples were added and incubated for 2 hours at room temperature, followed by another wash step. Subsequently, biotinylated detection antibodies for IL6 (BD Pharmingen, cat. #554546) or IL8 (Endogen, cat. #M-802-B) were applied for 1 hour, followed by incubation with Streptavidin-Conjugated Alkaline Phosphatase (BioRad, cat. #170-3554) for 30 minutes.

After extensive washing, PNPP substrate solution was added, and absorbance was measured at 405 nm with a reference wavelength of 690 nm using an iMark microplate reader (Bio-Rad). All samples were run in triplicate, and cytokine concentrations were calculated from standard curves with recombinant protein standards run in parallel.

### Western blotting

Cells were homogenized with lysis buffer (60 mM Tris-HCl pH 6.8, 2% sodium dodecyl sulfate, and a protease inhibitor cocktail (Cell Signaling Biotechnology, Beverly, MA)). Protein concentration was determined using the DC protein assay (Bio-Rad). 5-10 μg of total protein lysate per sample was separated via SDS-PAGE, transferred onto 0.45 μm Immobilon-PVDF membranes (Millipore, Billerica, MA), and blocked with 5% non-fat dry milk and 0.1% Tween 20 (BioRad) in 1X TBS and then incubated with antibodies against: phospho-AMPKα^Thr172^ (rabbit anti-AMPKα^Thr172^; cat number: 2535), total AMPKα (rabbit anti-AMPKα; cat number: 5831s), phospho-AKT (rabbit anti-pAKT (Ser473); cat number: 4060), total AKT (rabbit anti-AKT; cat number: 9272), Phospho-FoxO1a (rabbit anti-pFoxO1; cat number: 2599), total FoxO1a (rabbit anti-FoxO1; cat number: 2880), Phospho-NF-κB (rabbit anti-p NF-κB; cat number: 3033) and total NF-κB (rabbit anti-NF-κB; cat number: 4764). These antibodies were obtained from Cell Signaling Biotechnology (Danvers, MA). Tubulin was used as a loading control (anti-tubulin-hFAB RHODAMINE; cat number: 12004165, Bio-Rad). Anti-rabbit HRP-conjugated secondary antibodies were obtained from Jackson Immunoresearch (West Grove, PA). HRP signal was visualized using Immobilon Western chemiluminescent horseradish peroxidase substrate (Millipore, Billerica, MA). Chemiluminescent signals were captured using a Chemi-Doc MP Imaging System (Bio-Rad). Densitometric analysis was performed with Image Lab analysis software (Bio-Rad). Equal protein loading was confirmed using Mini-Protein TGX stain-free gels (Bio-Rad) visualized on the Chemi-Doc MP imaging system.

### Statistical analysis

All experiments were repeated at least three times using different strains of OFs with biological replicates (n=3) to drive the experiments. The data are expressed as the means ±SEM and were analyzed using Prism version 10.2.0 (GraphPad, Boston, MA). Student’s T-test and One-way analysis of variance (ANOVA) with multiple comparisons were used for statistical analysis. P-values of <0.05 were considered significant differences.

## Results

### TED orbital fibroblasts exhibit enhanced HA and cytokine responses to PDGFβ

Multiple growth factors and mediators activate OFs, leading to proliferation, HA production, and inflammation, which are critical in TED pathogenesis.^16,19,29,30^ We investigated the levels of secreted HA and inflammatory mediators in OFs in response to PDGFβ. OFs from non-TED and TED subjects were treated with PDGFβ (25 ng/mL) for 72 hours, and secreted HA levels were analyzed using agarose gel electrophoresis (**Figure 1A**). PDGFβ treatment induced high molecular weight HA synthesis in both non-TED and TED OFs, with TED OFs showing dramatically higher HA levels. Hyaluronidase treatment confirmed that the staining was due to HA and no other GAGs (**Figure S1A and S1B**). Quantification of HA using an ELISA further validated that TED OFs secrete significantly more HA, approximately 3-fold, than non-TED OFs in response to PDGFβ (**Figure 1B**). We also observed a significantly higher basal level of HA accumulation in TED OFs than non-TED OFs.

**Figure 1.**
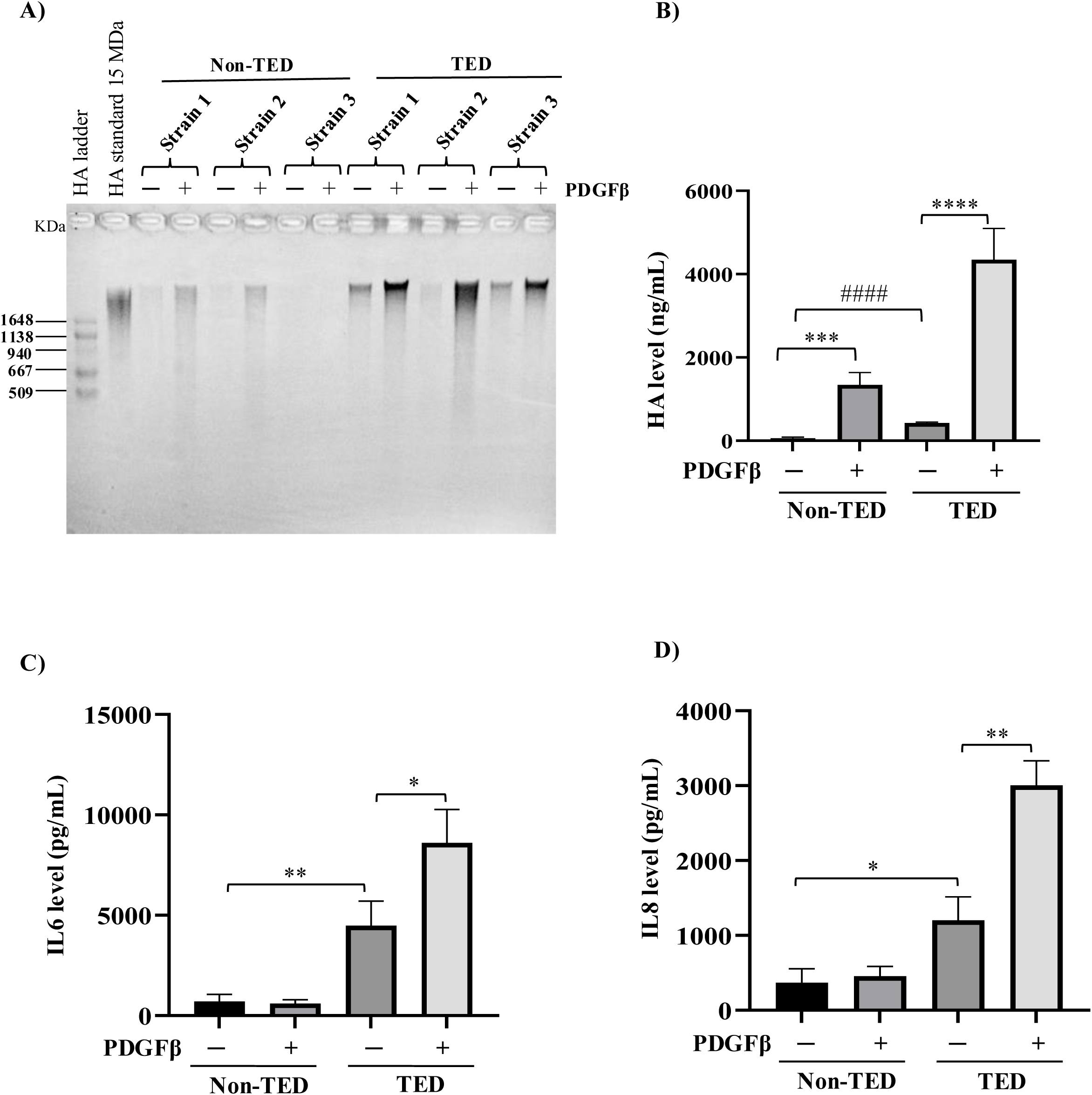
PDGFβ induces HA and pro-inflammatory cytokine production in both non-TED and TED OFs. A) non-TED (n=3) and TED (n=3) OFs were treated with or without PDGFβ for 72 hours. Secreted HA levels in media supernatant collected after treatment were analyzed on agarose gel electrophoresis and stained using Stains-All. 15 MDa and HiLadder were used as a size standard. B) Quantitative HA levels in both non-TED and TED after treatment were measured as described in the Methods section. C) and D) The relative production of IL6 and IL8 by OFs after PDGFβ treatment for 72 hours was measured with ELISA assay. The experiments were conducted in triplicate. The data represent mean ± SEM and were analyzed statistically using One-way ANOVA with Tukey’s multiple comparisons. Vehicle versus PDGFβ: \**P* ≤ 0.05, \*\**P* = 0.001, \*\*\**P <* 0.0008, \*\*\*\**P <* 0.0001, and non-TED versus TED: ^####^*P* ≤ 0.0001. HA, hyaluronan.

We next investigated whether the production of the inflammatory mediators, IL6 and IL8, were also elevated in TED OFs after PDGFβ treatment. IL6 and IL8 levels were analyzed by ELISA, and similarly to HA levels, we observed higher basal levels of IL6 and IL8 in TED OFs. PDGFβ treatment induced greater production of IL6 and IL8, by ∼14- and ∼6.5-fold, respectively, in the TED OFs compared to non-TED OFs (**Figure 1C and D**). Interestingly, in non-TED OFs, we did not see any induction of IL6 and IL8 in response to PDGFβ (**Figure 1C and D**). Our results suggest that TED OFs respond more robustly to PDGFβ, exhibiting higher levels of HA, IL6, and IL8 than non-TED OFs.

### PDGFβ suppresses AMPK activity in TED OFs

Prior studies have demonstrated that reduced activity of AMPK is associated with various diseases.^31,32^ We explored whether PDGFβ signaling affects AMPK activity in TED. To test this, TED OFs were treated with varying doses of PDGFβ (5-100 ng/mL) for 72 hours and AMPKα phosphorylation (phospho-AMPKα^Thr172^) was analyzed by western blot (**Figure 2A**). We observed a reduced AMPKα^Thr172^ phosphorylation level with increasing concentration of PDGFβ. At doses more than 10 ng/ml, a significant reduction in the phosphorylation level of AMPKα^Thr172^ was observed (**Figure 2B**). These data show that PDGFβ signaling reduces the phosphorylation level of AMPKα^Thr172^ and, thus, suppresses the activity of AMPK in TED OFs.

**Figure 2.**
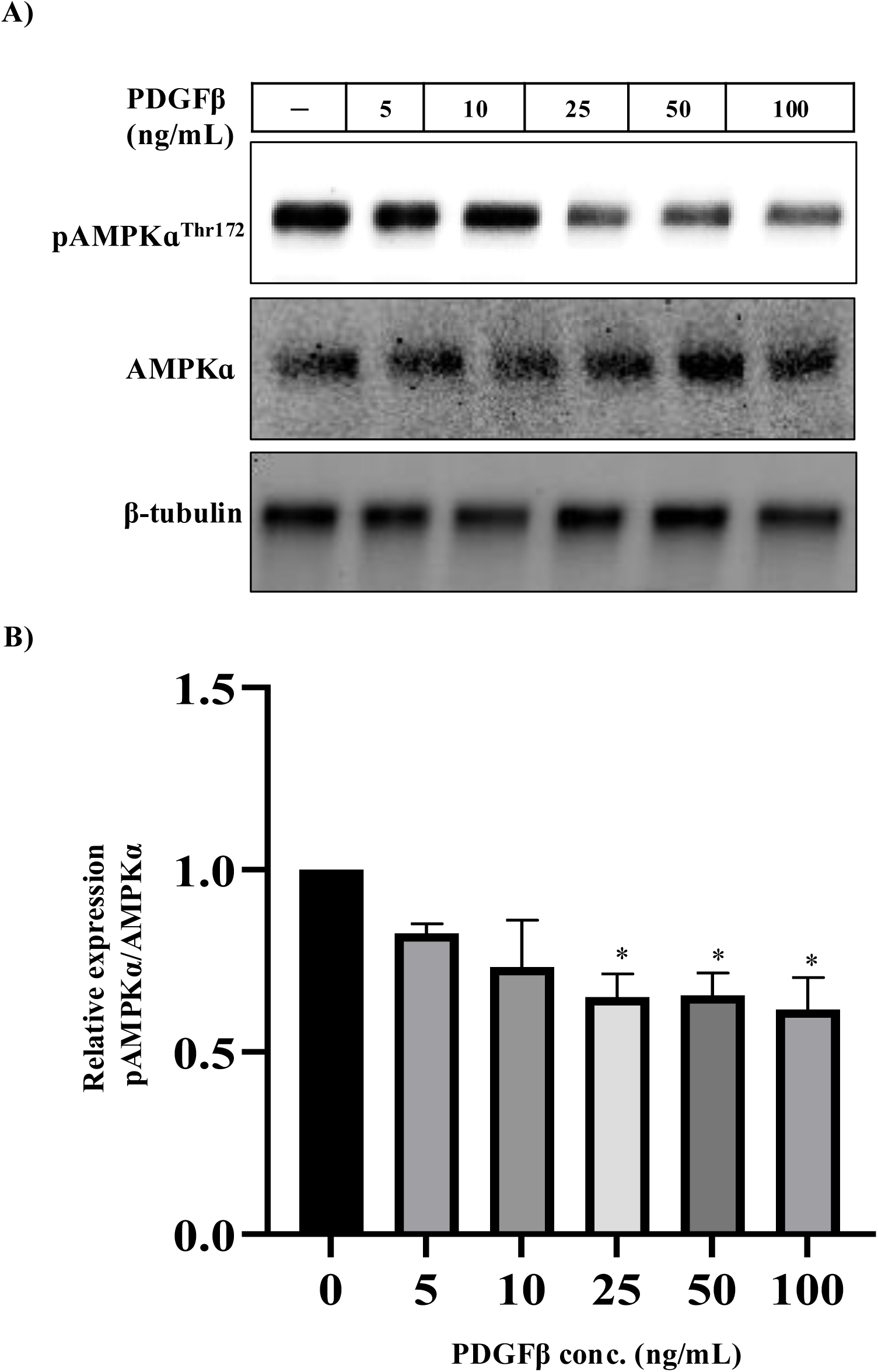
PDGFβ suppresses AMPK signaling in TED OFs. A) TED OFs were treated with different amounts of PDGFβ as indicated for 72 hours; thereafter, cells were analyzed by Western blot for pAMPKɑ^Thr172^, total AMPKɑ and β-tubulin, which served as a loading control. B) Relative expression level of pAMPKɑ^Thr172^ normalized to total AMPKɑ. Band intensities were quantified using Image Lab software. The experiment was repeated in three TED OF strains with representative results. Data represent mean ± SEM. One-way ANOVA with Sidak’s multiple comparisons was used to compare vehicle versus PDGFβ treatments. *p≤ 0.05.

### Metformin increases AMPK phosphorylation and decreases IL6 and IL8 levels

Metformin is an antidiabetic drug that indirectly activates AMPK by inhibiting mitochondrial complex I, leading to altered cellular energy metabolism and various metabolic effects.^33,34^ To examine the effect of metformin on AMPK phosphorylation, TED OFs were treated with different doses of metformin (10 to 5000 μM) for 72 hours and then analyzed by Western blot (**Figure 3A**). Metformin significantly increases AMPK phosphorylation (AMPKα^Thr172^) at 1000, 3000, and 5000 μM (**Figure 3B).** In TED patients, the severity of the disease is strongly associated with the presence of inflammatory mediators such as IL6 and IL8.^4^ Therefore, we next assessed the level of IL6 and IL8 in TED OFs following exposure to different doses of metformin. Metformin significantly reduced both IL6 and IL8 production in a dose-dependent manner (**Figure 3C and D**).

**Figure 3.**
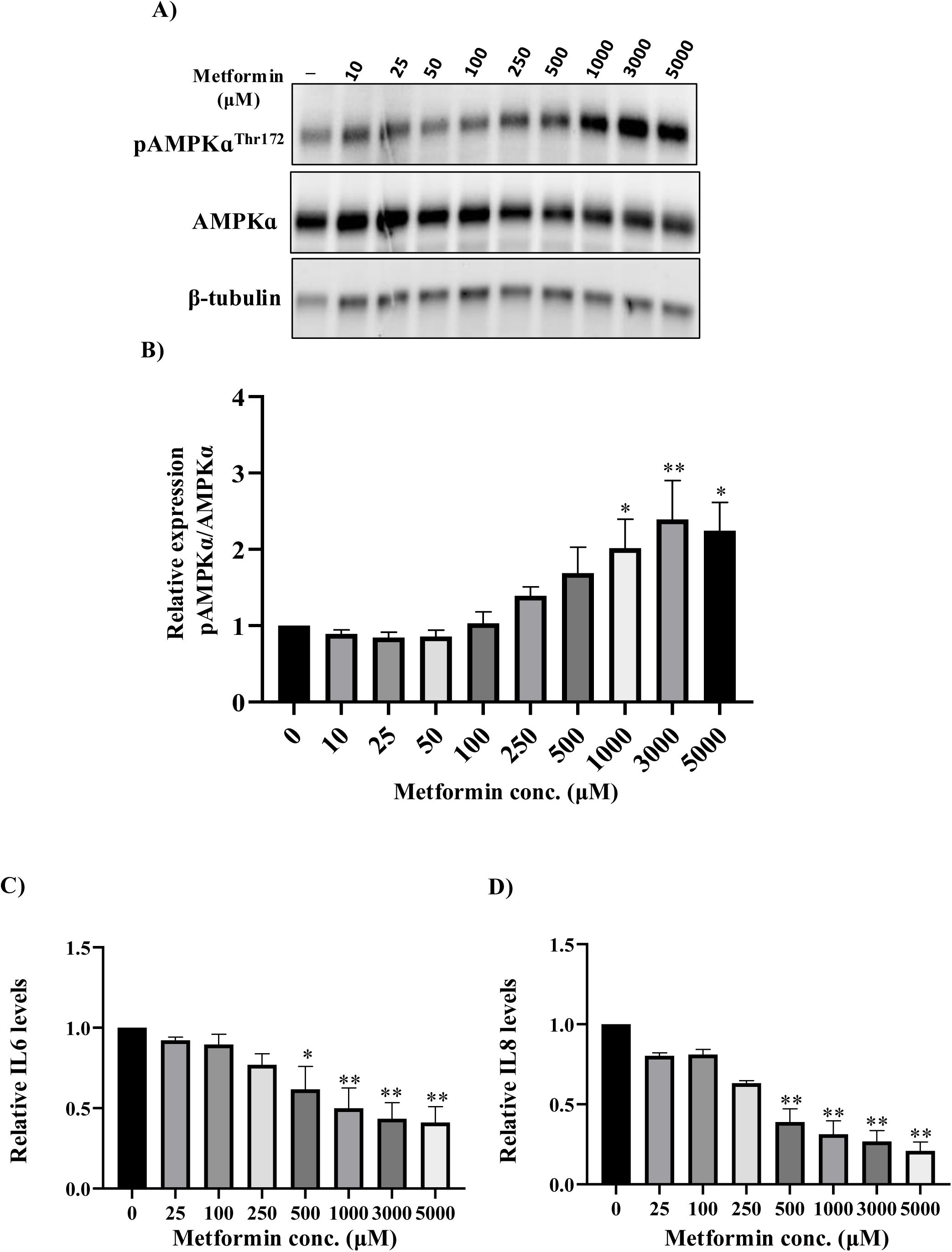
Metformin induces pAMPKɑ^Thr172^ level in TED OFs. A) TED OFs were treated with different doses of metformin as indicated for 72 hours; thereafter, cells were analyzed by Western blot for pAMPKɑ^Thr172^, total AMPKɑ, and β-tubulin (loading control). B) Relative levels of pAMPKɑ^Thr172^ normalized to total AMPKɑ in three strains with representative results. Band intensities were quantified using Image Lab software. C) and D) The relative production of IL6 and IL8 by OFs after metformin treatment for 72 hours was measured by ELISA. The experiments were conducted in triplicate on three different strains. The data represent mean ± SEM. One-way ANOVA with Dunnett’s multiple comparisons was used to compare vehicle versus metformin treatment. *p≤ 0.05, **p≤ 0.001.

### Metformin reduces PDGFβ-induced HA, IL6, and IL8 production in TED OFs

Since we saw that PDGFβ induced HA, IL6, and IL8 production in OFs (**Figure 1**) and that PDGFβ reduced AMPK activation (**Figure 2**), we next tested whether AMPK activation by metformin could attenuate PDGFβ signaling in TED OFs. We also tested AICAR, a direct activator of AMPK, on PDGFβ-induced responses. TED OFs were incubated with 1000 μM and 400 μM of metformin and AICAR, respectively, for 1 hour before PDGFβ was added for 72 hours. Cells were then harvested and analyzed by Western blot. Metformin and AICAR increased basal phospho-AMPKα^Thr172^ levels (F**igure 4A-B** **and S4**). Additionally, metformin and AICAR attenuated the PDGFβ-suppressed level of phospho-AMPKα^Thr172^ by ∼3-fold (**Figure 4A-B**). These data reveal a prominent role for AMPK activators in modulating PDGF signaling in TED OFs.

**Figure 4.**
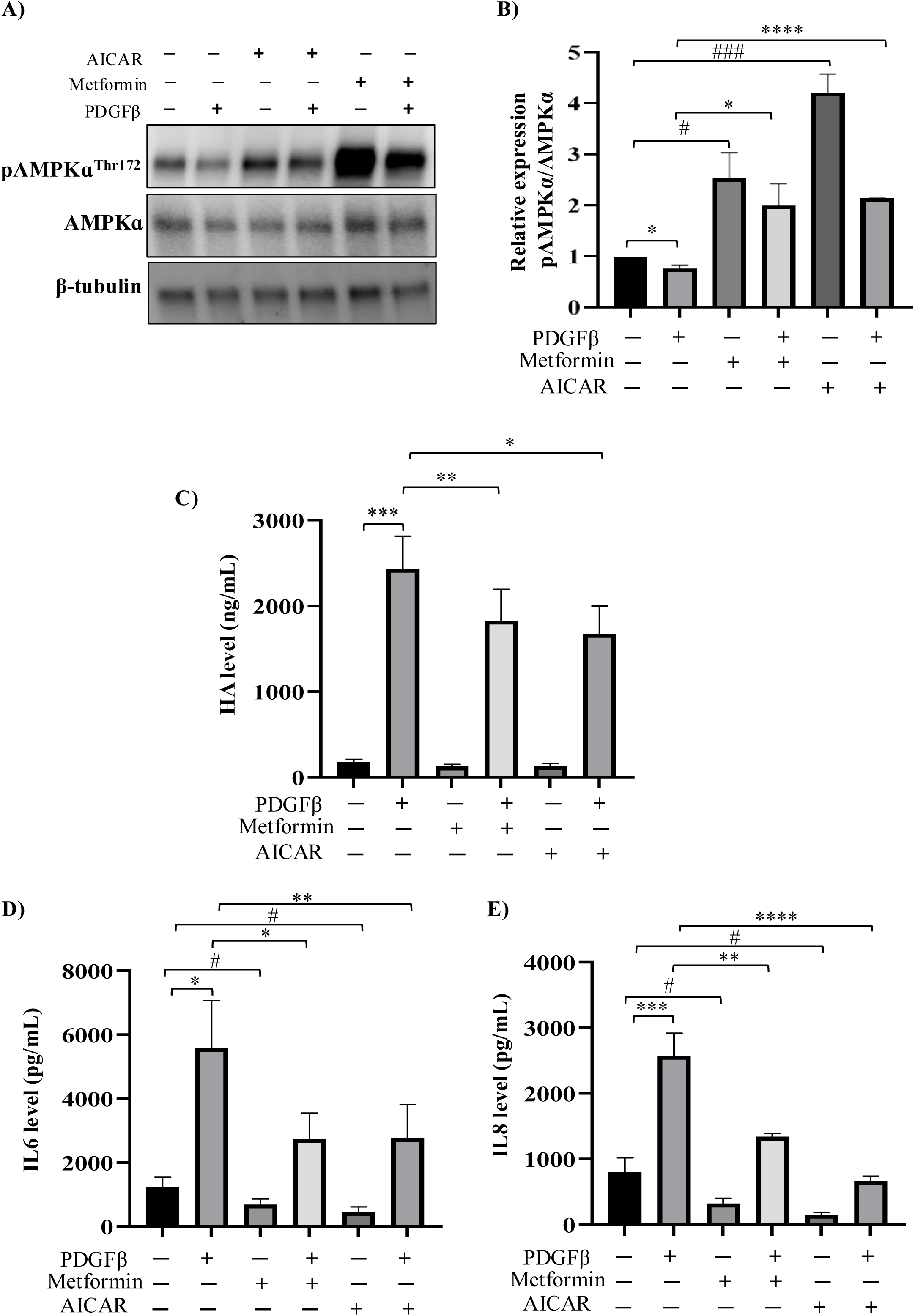
Metformin and AICAR attenuate PDGFβ-induced response in TED OFs. A) TED OFs were pretreated with AMPK activators metformin (1000 μM) or AICAR (400 μM) for 1 hour and then added PDGFβ (25 ng/mL) for the next 72 hours; thereafter, cells were analyzed by western blots for pAMPKɑ^Thr172^, total AMPKɑ and β-tubulin as a loading control. B) Relative levels of pAMPKɑ^Thr172^ normalized to total AMPKɑ in three strains with representative results. Band intensities were quantified using Image Lab software. C) Quantitative HA levels in TED OFs after treatment with PDGFβ and AMPK activators, metformin/AICAR, were measured with the HA ELISA assay. D) and E) The relative production of IL6 and IL8 by OFs after treatment with PDGFβ and AMPK activators, metformin/AICAR, was measured by ELISA. The experiments were conducted in triplicate on three different strains. The data represent mean ± SEM. Student’s T-test and One-way ANOVA with Tukey’s multiple comparisons were used to compare PDGFβ and PDGFβ + metformin/AICAR. *P ≤ 0.05, **P ≤ 0.001, ***P < 0.0009, ****P < 0.0001, and vehicle versus metformin/AICAR treatment. ^#^*P* ≤ 0.05, ^###^*P* ≤ 0.0005.

Prior studies have shown that when AMPK is activated, it can directly inhibit the synthesis of the enzyme Hyaluronan Synthase 2 (HAS2), which is involved in HA production.^35^ Given that activated AMPK can reduce the level of HA, we sought to determine if metformin and AICAR can suppress PDGFβ-induced production of HA. Interestingly, metformin and AICAR significantly decreased (∼1.4-fold) PDGFβ-induced HA production in TED OFs (**Figure 4C**). We further analyzed whether metformin and AICAR reduce PDGFβ-induced levels of IL6 and IL8. Remarkably, metformin suppressed PDGFβ-stimulated IL6 and IL8 levels by ∼2-fold, and in response to AICAR, this suppression was ∼2-fold (IL6) and ∼4-fold (IL8) (**Figure 4D-E**). Therefore, our data suggest that the AMPK activators metformin and AICAR suppress PDGFβ responses in TED OFs.

### Metformin suppresses PDGFβ-induced PI3K/AKT/FoxO1/NF-κB axis in TED OFs

We next sought to elucidate the mechanism(s) by which metformin attenuates PDGFβ responses in TED. Prior studies in OFs show that PDGFβ attributes its cellular response to the PI3K/AKT activation.^19^ Active AKT is key in regulating HA synthesis by increasing enzyme expression, such as HAS2.^36^ AKT also acts as a signaling molecule to promote cellular proliferation and inflammatory responses by activating NF-κB. Thus, we first sought to measure the phosphorylation of AKT at serine 473 (pAKT), which indicates active PI3K/AKT signaling. Metformin pretreatment significantly reduces (∼1.7-fold) PDGFβ-induced phosphorylation level of AKT, while AICAR did not show any significant changes in the level of pAKT (**Figure 5A-B and S5A**). As AICAR is a direct activator of AMPK, it is possible that metformin, for which some AMPK-independent roles are also identified, could be reducing pAKT level in an AMPK-independent manner.

**Figure 5.**
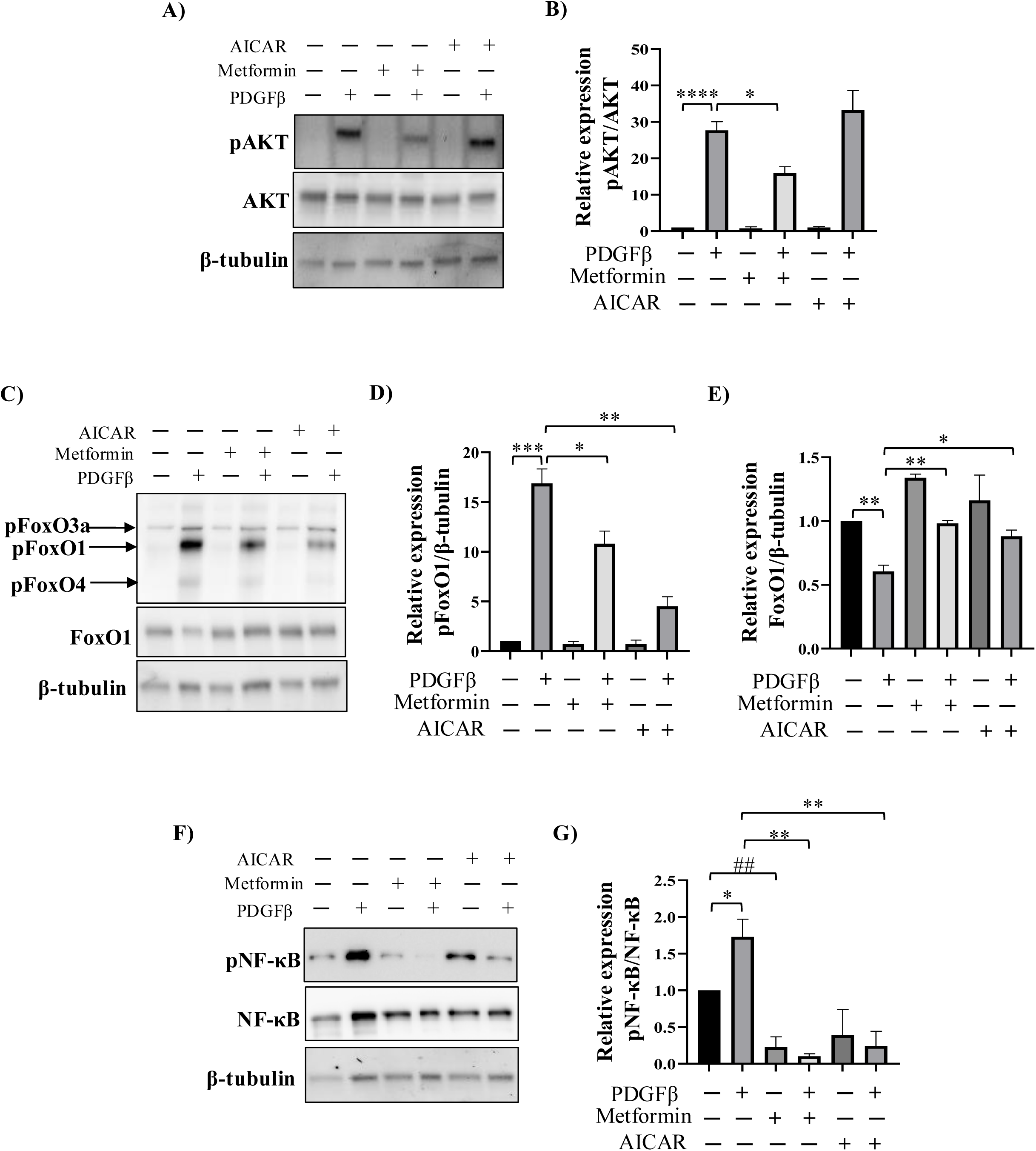
Metformin reduces PDGFβ-induced phosphorylation of Akt and FoxO1 in TED OFs. A-G) TED OFs were pretreated with AMPK activators metformin (1000 μM) or AICAR (400 μM) for 1 hour and then added PDGFβ (25 ng/mL) for 72 hours; thereafter, cells were lysed and analyzed by Western blot for pAkt, total Akt, pFoxO1, total FoxO, pNF-κB and total NF-κB, and β-tubulin as a loading control. Relative levels of pAkt were normalized to total Akt, pFoxo1 and total FoxO1 were normalized to β-tubulin, and pNF-κB was normalized to total NF-κB in three strains with representative results shown. Band intensities were quantified using Image Lab software. The data represent mean ± SEM. Student’s T-test and One-way ANOVA with Dunnett’s multiple comparisons were used to compare PDGFβ and PDGFβ + metformin/AICAR. *P ≤ 0.05, **P ≤ 0.009, ***P < 0.0002 ****P < 0.0001 and vehicle versus metformin/AICAR treatment. ^##^*P* ≤ 0.005.

To further pursue the signaling pathway, we investigated the expression and phosphorylation status of FoxO1. AMPK has been shown to enhance the transcriptional activity of FOXOs.^37^ Activated AKT phosphorylates FoxO1 at serine residue 24 and acts as a negative regulator, blocking nuclear transport and thus transcriptional activity. FoxO1 plays a central role in regulating cell growth, survival, and metabolic homeostasis downstream of AMPK and AKT signaling.^38–40^ Thus, we measured the expression of the phosphoSer24-FoxO1 (pFoxO1) in TED OFs. PDGFβ treatment increased pFoxO1, and metformin and AICAR suppressed pFoxO1 by ∼2.5-fold and ∼5.4-fold, respectively (**Figure 5C-D and S5B**). We also observed that PDGFβ treatment reduces basal level expression of FoxO1, and addition of metformin and AICAR restores PDGFβ-reduced levels of FoxO1 (**Figure 5C, E, and SB**), which is consistent with previous studies demonstrating that PDGF induces rapid phosphorylation and then degradation of FoxO1 protein by the proteasome.^39^ These data suggest that metformin inhibits the phosphorylation of pFoxO1 in an AMPK-dependent manner.

To further define the signaling pathway affected by PDGFβ, we examined the levels and activation status of nuclear factor-κB (NF-κB). NF-κB plays a central role in inflammation by acting as a transcription factor that promotes the expression of pro-inflammatory genes, including IL6 and IL8. AKT and FoxO1 also modulate NF-κB activity, forming an integrated signaling network that governs inflammatory responses.^41,42^ To assess NF-κB activation, we measured phosphorylation of the p65 subunit at serine 536 (p-NF-κB Ser536), a marker of transcriptional activation that facilitates nuclear translocation and gene expression.^43^ Metformin and AICAR significantly reduced PDGFβ-induced phospho-NF-κB levels by ∼17-fold and ∼8.5-fold in TED OFs (**Figure 5F-G and S5C**). These findings suggest that metformin suppresses PDGFβ-driven inflammation by targeting the PI3K/AKT/FoxO1/NF-κB axis in orbital fibroblasts.

## DISCUSSION

TED is driven by complex, multifaceted pathophysiological processes that involve excessive HA accumulation, inflammatory cytokine production, and tissue remodeling within the orbit.^1,3^ OFs are central mediators of these events, yet therapeutic strategies directly targeting fibroblast pathogenicity remain limited. Here, we provide novel evidence that PDGFβ signaling suppresses AMPK activity in TED OFs, promoting pathological HA synthesis and inflammatory cytokine production. Furthermore, we demonstrate that AMPK activation by metformin can counter PDGFβ signaling, identifying a new avenue for therapeutic intervention in TED. Activation of AMPK attenuates PDGFβ-induced responses by blocking the PI3K/AKT/FoxO signaling network in TED OFs (**Figure 6).**

**Figure 6.**
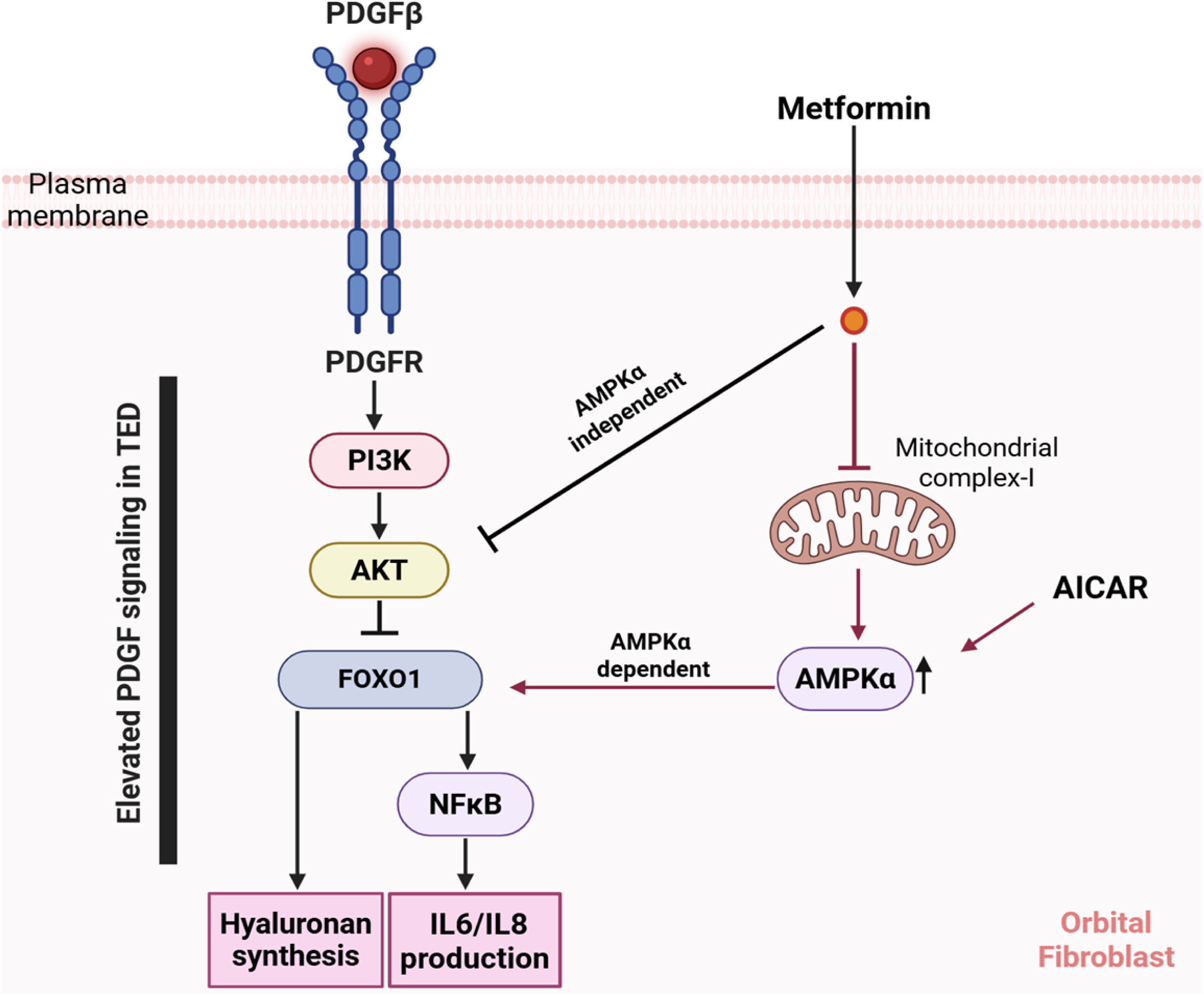
Model depicting AMPK activator metformin attenuates PDGFβ-induced PI3K/AKT/FoxO1/NF-κB network in TED. PDGFβ activates PI3K/pAKT/FoxO1/NF-κB signaling cascades in TED OFs by phosphorylating AKT. The phosphorylated form of Akt then inactivates FoxO1, a transcription factor, by phosphorylating it at serine 24. Both AKT and FoxO1 are involved in inducing the synthesis of HA and producing inflammatory mediators such as IL6 and IL8 through NF-κB. Metformin inhibits the PI3K/AKT/FoxO1/NF-κB pathway components through both AMPKα-dependent and independent mechanisms. At the same time, AICAR, as opposed to metformin, works primarily through an AMPKα-dependent mechanism to attenuate PDGFβ responses.

Our findings reveal that TED OFs display a heightened sensitivity to PDGFβ stimulation compared to non-TED OFs, manifesting in markedly increased HA, IL6 and IL8 production. PDGFβ levels are significantly higher in orbital tissue from TED patients compared to non-TED subjects.^43–46^ This intrinsic hyperresponsiveness likely reflects disease-associated alterations in fibroblast signaling states acquired during TED pathogenesis.^16,19,47^

Further, we demonstrate that metformin increases the AMPK activity and reduces the production of basal levels of inflammatory cytokines IL6 and IL8 in TED OFs, which has been reported in TED and other autoimmune diseases.^21,24,25^ We propose that metformin, by its ability to activate AMPK, can block pathogenic PDGFβ signaling. PDGFβ induces HA production dramatically in TED OFs, and elevated HA is a hallmark of TED. Suppression of AMPK phosphorylation by PDGFβ may represent a critical pathological mechanism underlying this fibroblast activation, aligning with broader evidence linking impaired AMPK signaling to fibrosis, inflammation, and metabolic dysfunction across multiple diseases.^21,22,24–27^

Importantly, we show that restoring AMPK activity with metformin reverses PDGFβ-mediated AMPK suppression and significantly inhibits downstream pathways. Metformin reduced HA accumulation and inflammatory cytokine production under basal conditions and after PDGFβ stimulation. Mechanistically, metformin suppressed PI3K/AKT signaling and reduced inhibitory phosphorylation of FoxO1, a key transcription factor involved in extracellular matrix regulation and inflammation.^48^

Our results suggest that metformin modulates orbital fibroblast responses in TED through both AMPK-dependent and AMPK-independent mechanisms. Prior studies have implicated the PI3K/AKT pathway in promoting HA synthesis and inflammation in orbital fibroblasts, often downstream of IGF-1R or TSHR activation. Crosstalk between these receptors has been proposed as a key driver of disease pathogenesis, enhancing fibroblast activation and proliferation.^13,30^ Analogous to observations in pancreatic ductal adenocarcinoma, where metformin-induced AMPK activation disrupted IGF-1R and GPCR crosstalk and inhibited mTORC1-dependent tumor cell proliferation^49^, a similar mechanism may be operative in TED. Metformin may interfere with pathogenic cooperation between TSHR and IGF-1R via metabolic reprogramming effects; however, further studies are needed to confirm this potential regulatory axis in TED OFs. Future experiments using activation of TSHR and IGF-1R in OFs, with or without metformin treatment, could help clarify whether metformin disrupts this cooperative signaling to attenuate fibroblast activation.

Our findings also extend this mechanistic paradigm to PDGF signaling and identify AMPK as a key modulator of both fibrotic and inflammatory responses in TED. Given metformin’s well-established safety profile, affordability, and global clinical use, these results support its potential for therapeutic repositioning in TED. This approach may be particularly beneficial during the early, active phase of disease, when fibroblast activation and tissue remodeling are predominant. However, metformin may also offer therapeutic value in chronic-stage TED by limiting the excessive accumulation of HA that contributes to orbital congestion and enlargement.

Metformin is the most widely prescribed drug for the treatment of type 2 diabetes mellitus (T2DM), the most common endocrine disorder, associated with significant morbidity and mortality.^55^ Epidemiological studies have reported a higher frequency and severity of TED in patients with coexisting T2DM.^50,51^ However, the mechanistic connection between T2DM and TED remains incompletely understood. One potential link involves insulin resistance—a hallmark of T2DM—which promotes dysregulated lipid metabolism and hepatic fat accumulation.^52^ Insulin resistance also reduces levels of IGF-1 binding proteins, leading to elevated free IGF-1.^53^ Increased IGF-1 bioavailability may contribute to orbital tissue remodeling by activating IGF-1R signaling pathways, which are well established in TED pathogenesis. Inhibitors of IGF-1R signaling, such as teprotumumab, have shown efficacy in reducing adipogenesis and HA production in TED, although their use may be limited by adverse effects, including hearing loss, muscle spasms, and hyperglycemia. Notably, metformin could potentially mitigate teprotumumab-induced hyperglycemia if used in combination therapy, while also exerting direct anti-fibrotic and anti-inflammatory effects in TED.

Previous *in vitro* studies have demonstrated that TSHR and IGF-1R co-activation in orbital fibroblasts signals through the PI3K/AKT pathway to drive pathological remodeling. Our current findings extend this framework by showing that PDGF signaling, also elevated in TED orbital tissue, can be attenuated by metformin via AMPK activation and downstream inhibition of the PI3K/AKT pathway. This suggests that metformin may disrupt multiple converging pro-pathogenic signaling networks in TED. Given the mechanistic parallels between T2DM and TED, our results support the potential for metformin to serve as a repurposed therapeutic agent in TED. Notably, no clinical studies to date have evaluated whether anti-diabetic therapies, such as metformin, influence TED onset or progression. It would therefore be of great interest to examine whether diabetic patients treated with metformin exhibit a lower incidence or reduced severity of TED compared to those receiving alternative anti-diabetic treatments.

In summary, this study identifies a previously unrecognized role for PDGFβ-mediated suppression of AMPK in promoting pathological fibroblast activation in TED. By restoring AMPK activity, metformin attenuates PDGFβ-induced HA, IL6, and IL8 production, revealing a novel mechanism through which metformin may exert therapeutic benefit. These findings expand the therapeutic landscape for TED, suggesting that targeting fibroblast signaling pathways beyond IGF-1R and TSHR, particularly via modulation of metabolic regulators such as AMPK, could provide new strategies to limit disease progression. Given its broad clinical use, favorable safety profile, and affordability, metformin emerges as a promising candidate for future clinical trials aimed at preventing or reversing early orbital tissue remodeling in TED.

## FIGURE LEGENDS

**Figure S1.**
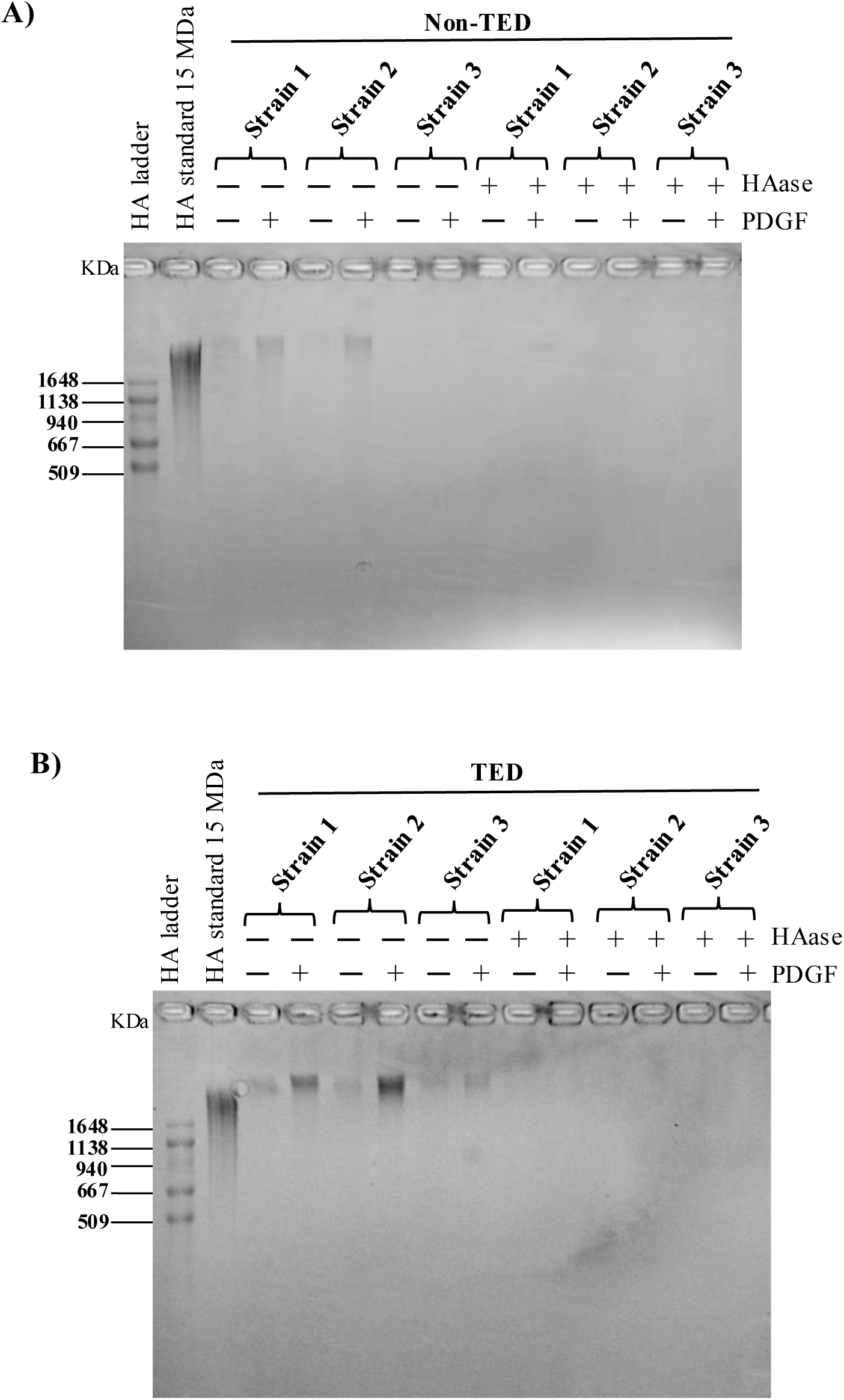
Treatment with hyaluronidase removes all staining in HA agarose gels. To confirm the staining was HA specific, aliquots of the samples were treated with 1 U/mL of Hyaluronidase and incubated overnight at 37°C. The following day, samples were processed and separated on an agarose gel. Hyaluronidase treatment confirmed that the staining was due to HA and not other GAGs.

**Figure S2.**
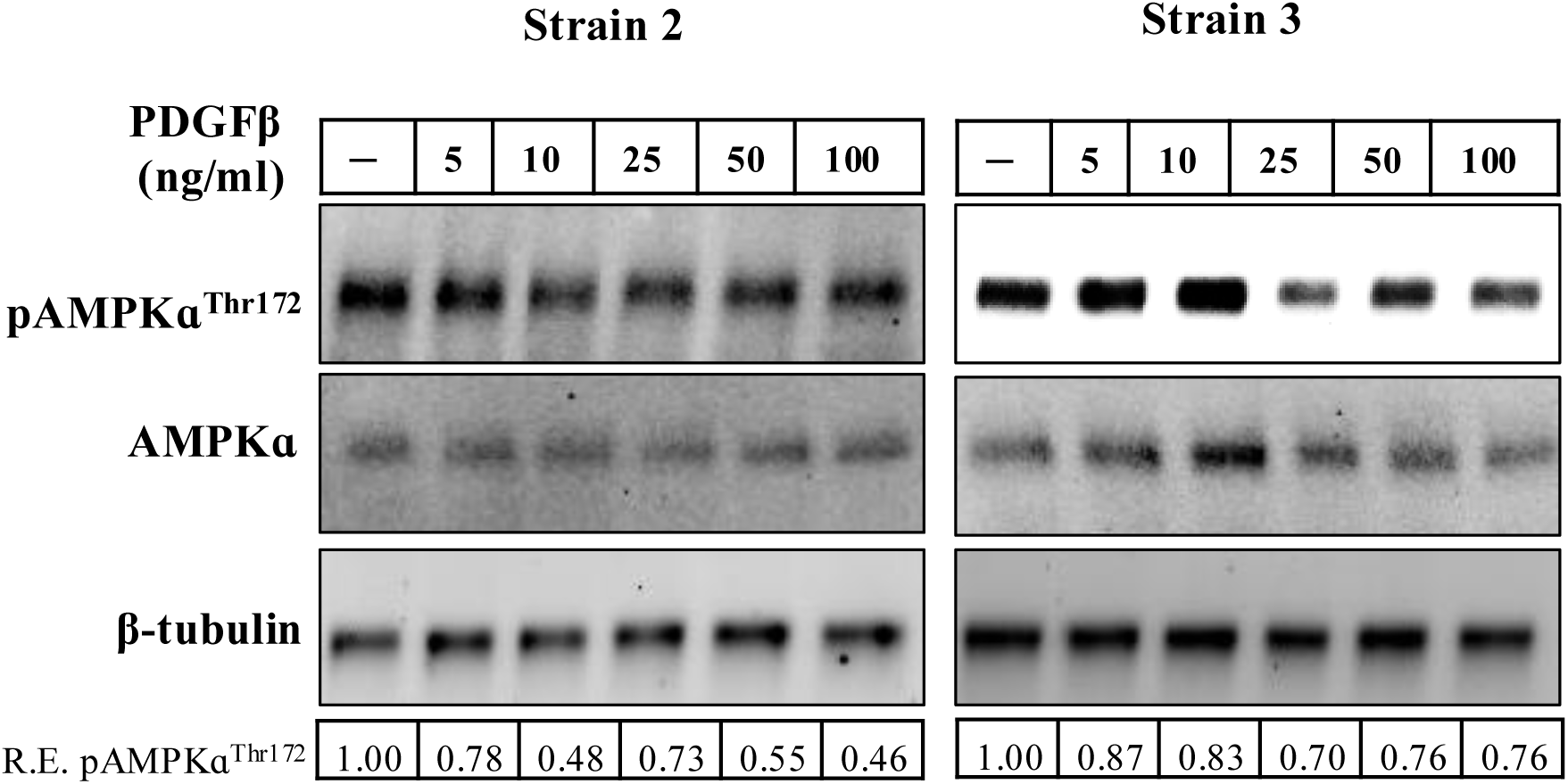
PDGFβ suppresses AMPK signaling in TED OFs. A) TED OFs were treated with different doses of PDGFβ as indicated for 72 hours; thereafter cells were analyzed by western blots for the expression of pAMPKɑ^Thr172^, total AMPKɑ and β-tubulin which served as a loading control.

**Figure S3.**
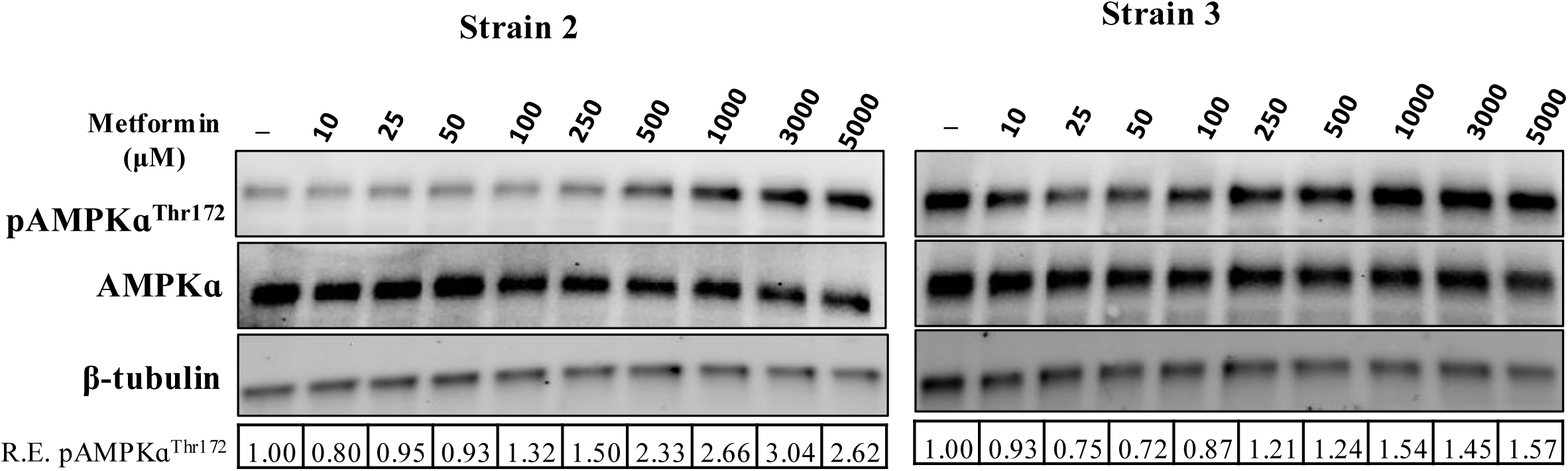
Metformin induces pAMPKɑ^Thr172^ level in TED OF. A) TED OF were treated with different doses of metformin as indicated for 72 hours; thereafter, cells were analyzed by western blots for pAMPKɑ^Thr172^, total AMPKɑ and β-tubulin (loading control).

**Figure S4.**
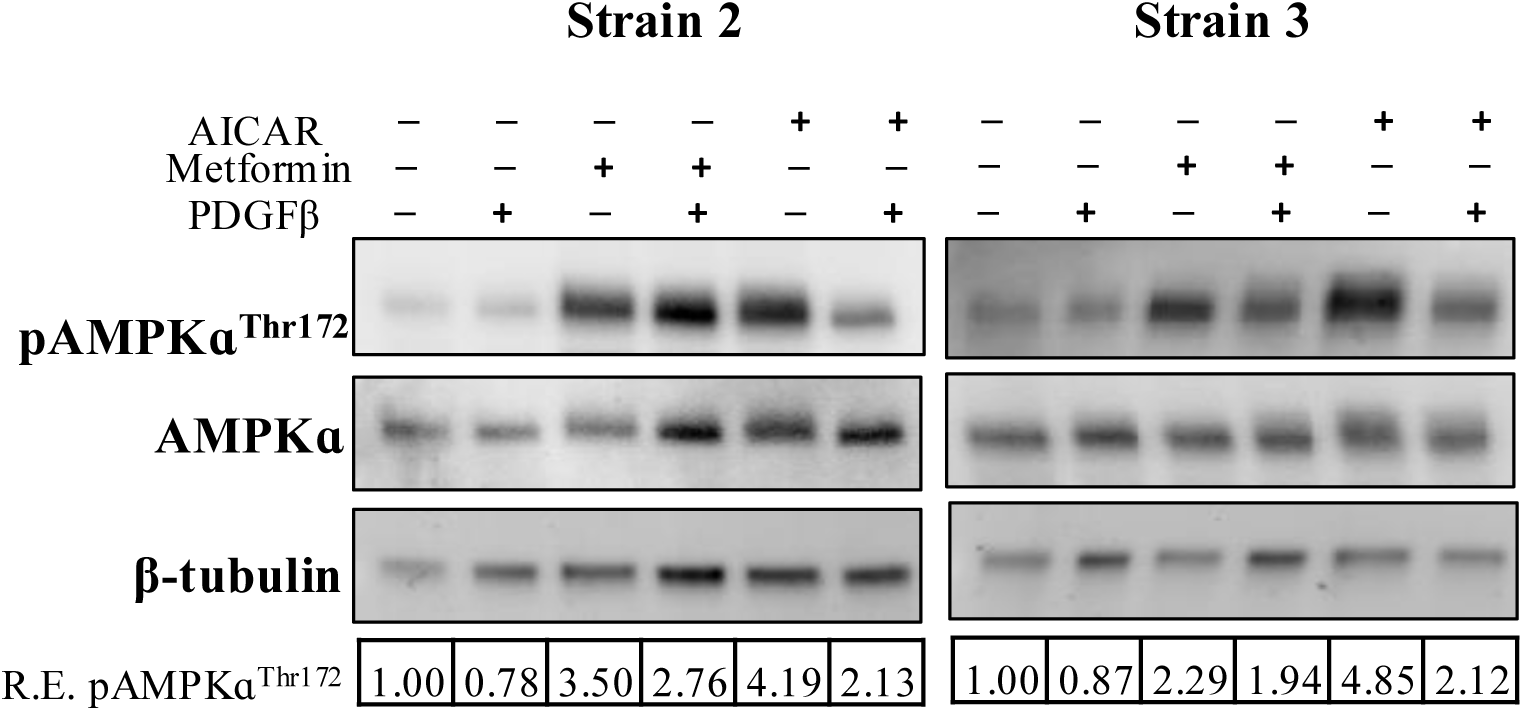
Metformin and AICAR attenuates PDGFβ induced response in TED OFs. TED OFs were pretreated with AMPK activators metformin (1000 μM) or AICAR (400 μM) for 1 hour and then added PDGFβ (25 ng/ml) for the next 72 hours; thereafter cells were analyzed by western blots for pAMPKɑ^Thr172^, total AMPKɑ and β-tubulin as a loading control.

**Figure S5.**
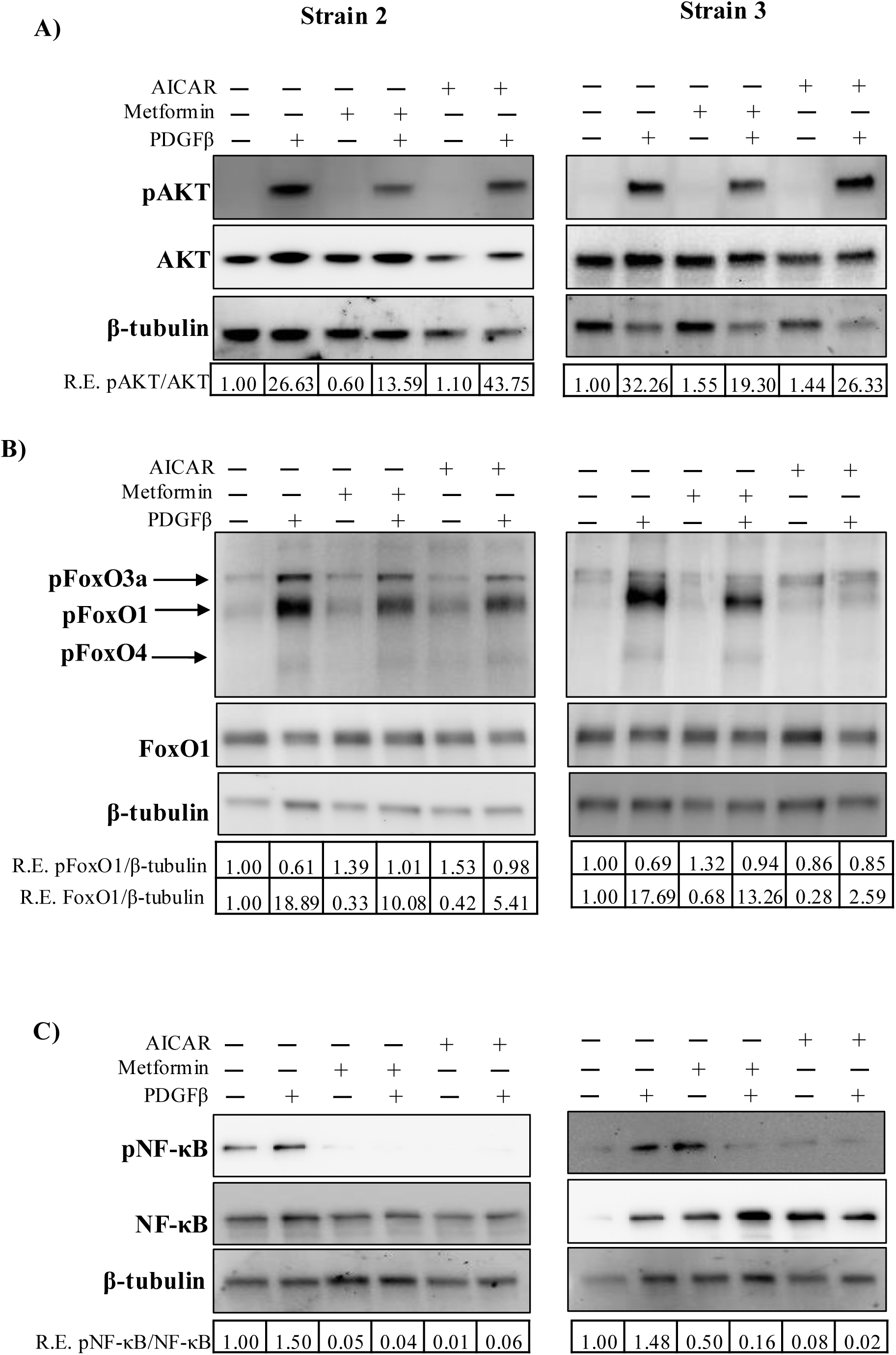
Metformin inhibits PDGFβ induced PI3K/AKT signaling in TED OFs. **A), B) and C)** TED OFs were pretreated with AMPK activators metformin (1000 μM) or AICAR (400 μM) for 1 hour and then added PDGFβ (25 ng/ml) for 72 hours; thereafter cells were lysed and analyzed by western blots for pAKT, total AKT, pFoxo3a, total Foxo3a, pNFκB and total NFκB and β-tubulin as a loading control.

